# *Phodopus roborovskii* SH101 as a systemic infection model of SARS-CoV-2

**DOI:** 10.1101/2021.03.10.434891

**Authors:** Chongkai Zhai, Mingda Wang, Hea-Jong Chung, Md. Mehedi Hassan, Seungkoo Lee, Hyeon-Jin Kim, Seong-Tshool Hong

**Affiliations:** Department of Biomedical Sciences and Institute for Medical Science, Jeonbuk National University Medical School, Jeonju, Jeonbuk 54907, South Korea; Gwangju Center, Korea Basic Science Institute, Gwangju 61751; Department of Anatomic Pathology, School of Medicine, Kangwon National University, Kangwon National University Hospital, 1 Gangwondaehak-gil, Chuncheon, Gangwon 24341, South Korea; JINIS BDRD Institute, JINIS Biopharmaceuticals Inc., 224 Wanjusandan-6-ro, Bongdong, Jeonbuk 55315, South Korea

## Abstract

Severe acute respiratory syndrome CoV-2 (SARS-CoV-2) is currently causing a worldwide threat with its unusually high transmission rates and rapid evolution into diverse strains. Unlike typical respiratory viruses, SARS-CoV-2 frequently causes systemic infection by breaking the boundaries of the respiratory systems. The development of animal models recapitulating the clinical manifestations of COVID-19 is of utmost importance not only for the development of vaccines and antivirals but also for understanding the pathogenesis. However, there has not been developed an animal model for systemic infection of SARS-CoV-2 representing most aspects of the clinical manifestations of COVID-19 with systemic symptoms. Here we report that a hamster strain of *Phodopus roborovskii* SH101, a laboratory inbred hamster strain of *P. roborovskii,* displayed most symptoms of systemic infection upon SARS-CoV-2 infection as in the case of the human counterpart, unlike current COVID-19 animal models. *P. roborovskii* SH101 post-infection of SARS-CoV-2 represented most clinical symptoms of COVID-19 such as snuffling, dyspnea, cough, labored breathing, hunched posture, progressive weight loss, and ruffled fur, in addition to high fever following shaking chills. Histological examinations also revealed a serious right-predominated pneumonia as well as slight organ damages in the brain and liver, manifesting systemic COVID-19 cases. Considering the merit of a small animal as well as its clinical manifestations of SARS-CoV-2 infection in human, this hamster model seems to provide an ideal tool to investigate COVID-19.

**Author summary:** Although the current animal models supported SARS-CoV-2 replication and displayed varying degrees of illness after SARS-CoV-2 infection, the infections of SARS-CoV-2 were mainly limited to the respiratory systems of these animals, including hACE2 transgenic mice, hamsters, ferrets, fruit bats, guinea pigs, African green monkey, Rhesus macaques, and Cynomolgus macaques. While these animal models can be a modest model for the respiratory infection, there is a clear limit for use them in the study of COVID-19 that also displays multiple systemic symptoms. Therefore, the development of an animal model recapitulating COVID-19-specific symptoms such as the right-predominated pneumonia would be the utmost need to overcome the imminent threat posed by COVID-19. We identified a very interesting hamster strain, *Phodopus roborovskii* SH101, which mimics almost all aspects of the clinical manifestations of COVID-19 upon SARS-CoV-2 infection. Unlike the current animal models, SARS-CoV-2-infected *P. roborovskii* SH101 not only displayed the symptoms of respiratory infection but also clinical manifestations specific to human COVID-19 such as high fever following shaking chills, serious right-predominated pneumonia, and minor organ damages in the brain and liver.

## Introduction

The emergence of COVID-19 upon the infection of SARS-CoV-2 in 2019 poses a mounting threat to the world. In COVID-19, the infection of SARS-CoV-2 in the respiratory system causes fever, shaking chills, headache, and fatigue followed by respiratory symptoms such as cough, sneeze, sore throat, chest pain, atypical pneumonia, *etc* [1,2]. The clinical features of COVID-19, however, are not limited to the consequences of a respiratory infection [3]. Unlike a typical respiratory infection, SARS-CoV-2 infects the organs other than the respiratory system from the beginning or subsequent to respiratory infection, leading to diarrhea, loss of sense of smell or taste, neuroinflammation manifested as encephalitis, meningitis, acute cerebrovascular disease, and Guillain Barré Syndrome (GBS), multisystem inflammatory syndrome, *etc* [4–6]. More interestingly, atypical pneumonia by SARS-CoV-2 is observed with heavy predominance in the right lung [7]. The right-over-left predominance in pneumonia is one of the hallmarks of COVID-19 disease [8,9]. As the consequences of these systemic infection, patients recovered from severe or even mild COVID-19 are frequently suffering from fatigue, heart palpitations, changes in lung function, muscle weakness, memory loss, concentration, brain fog, depression, anxiety, *etc* [10–12].

The mutation rate in RNA virus such as SARS-CoV-2 is dramatically high, up to a thousand times higher than that of DNA virus, contributing to rapid evolution into its variants. Considering the nature of RNA genome and the number of already happened cases of outbreak, the emergence of new strains from furtively circulating SARS-CoV-2 even after herd immunity by vaccination seems to be inevitable. In fact, the world is already observing the emergence of multiple variants of SARS-CoV-2. Given the number of already spread outbreak cases, an appropriate surrogate for human COVID-19 is needed to overcome the imminent threats of COVID-19 from a future emergence of variant strains. Moreover, animal models could help shed light on important aspects of human COVID-19 in ways that are not easily addressed or feasible in humans, such as how SARS-CoV-2 causes systemic infection.

After the outbreak of COVID-19, various animal models for COVID-19 have been developed. The current animal models, such as hACE2 transgenic mice, hamsters, ferrets, fruit bats, guinea pigs, African green monkey, Rhesus macaques, and Cynomolgus macaques, supported SARS-CoV-2 replication and displayed varying degrees of illness when the virus was delivered into the respiratory tract of these animals [13–17]. However, the infections of SARS-CoV-2 in these animals were mainly limited to the respiratory systems. Since these animal models mostly recapitulated the infections of the respiratory system, each of these animals has a limited utility in the study of COVID-19 [18–22]. Therefore, the development of an animal model recapitulating systemic infection, including COVID-19-specific facet such as the right-predominated pneumonia, would be the utmost need to overcome the imminent threat posed by COVID-19.

In this work, we identified a laboratory-inbred strain of *Phodopus roborovskii*, *P. roborovskii* SH101, representing the systemic infection of human COVID-19. In addition to systemic infection, the infection of *P. roborovskii* SH101 with SARS-CoV-2 caused almost all clinical symptoms represented by human COVID-19, including high fever, shaking chills, serious right-predominated pneumonia, and the viral distribution pattern as well as evident respiratory and behavioral symptoms.

## Results

### A hamster strain recapitulating COVID-19 after SARS-CoV-2 infection was identified

To develop an ideal animal model of COVID-19, we screened several hundred strains of mice, rats, guinea pigs, and hamsters, and identified a hamster strain, *Phodopus roborovskii* SH101, which was highly sensitive to SARS-CoV-2 infection. *P. roborovskii* SH101 (abbreviated as SH101) is a laboratory-inbred hamster strain with prominent white patches above eyes and at the base of ears (Fig 1A, B). The average adult body weights of SH101 were about 21 g for males and 20 g for females, respectively.

**Fig 1.**
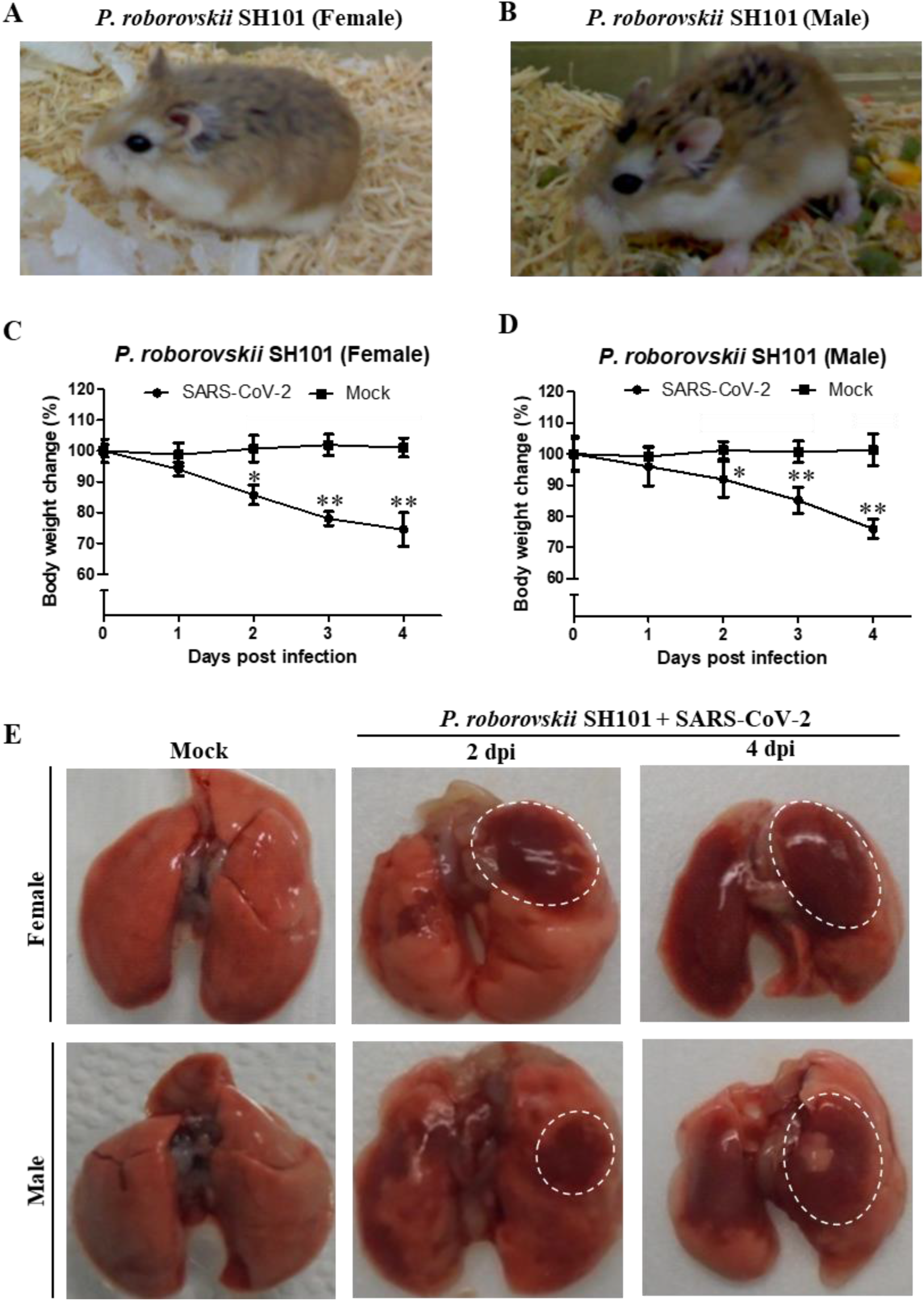
Identification of a small animal model for SARS-CoV-2 infection representing most clinical features of COVID-19. (A, B) The photographic image of adult female (A) and male (B) *Phodopus roborovskii* SH101, a laboratory inbred strain. (C, D) The body weight changes for female (C) and male (D) *P. roborovskii* SH101 post-infection of SARS-CoV-2. The body weights were measured daily for 5 days (up to 4 dpi) (n = 6). Data are presented as mean ± SD. The statistical significances are marked on the graphs as * *P* < 0.05 and ** *P* < 0.01. (E) The photographic images of the dissected lungs of the SARS-CoV-2-infected hamster with right-predominant pneumonia indicated as white dotted circles.

As in the case of COVID-19, respiratory symptoms were immediately noticed in the SH101 hamsters infected with SARS-CoV-2 (Video S1~S4). The hamsters showed clear signs of respiratory symptoms such as snuffling, dyspnea, cough, labored breathing, ruffled fur, and sneeze. Along with the respiratory symptoms, the very active behavior otherwise typical for the hamsters was dramatically reduced while hunched posture was observed starting from 1-day post-infection (dpi). Other than these clinical manifestations associated with the severe respiratory and systemic infection, some hamsters showed shaking chills (Video S3) after 2 dpi which resembled the shaking chills of human patients of COVID-19. Also, progressive and significant weight loss had been observed from 2 to 4 dpi (Fig 1C, D). All individuals were terminally ill, and the mortality rate of the SH101 hamsters was 83% by 4 dpi.

Most interestingly, a unique uneven distribution of pneumonia was noticed immediately by gross examination of the lung specimens (Fig 1E). The dissected lungs of the infected SH101 hamsters showed the right-predominated pneumonia just like as in the case of COVID-19. It was remarkably interesting to note the right-predominated pneumonia of the hamster because one of the most peculiar clinical manifestations of COVID-19 is the right-over-left predominated pneumonia [23–25].

### *P. roborovskii* SH101 infected with SARS-CoV-2 progressed similarly with human COVID-19

Although the induction of fever in COVID-19 is an essential hallmark of SARS-CoV-2 infection, none of the current animal models, including primates, showed an induced fever by SARS-CoV-2 infection. In this study, the body temperature measurement by an infrared thermographic method revealed the induction of fever in SH101 hamsters immediately after infection of SARS-CoV-2 (Fig 2). The elevated body temperatures of the infected SH101 hamsters were sharply dropped as the infection progressed to the terminal stage. Being one of the cardinal features in COVID-19 [26,27], the fever induction after SARS-CoV-2 infection seems to indicate that the SH101 hamsters simulate COVID-19 symptoms most closely among animal models of COVID-19.

**Fig 2.**
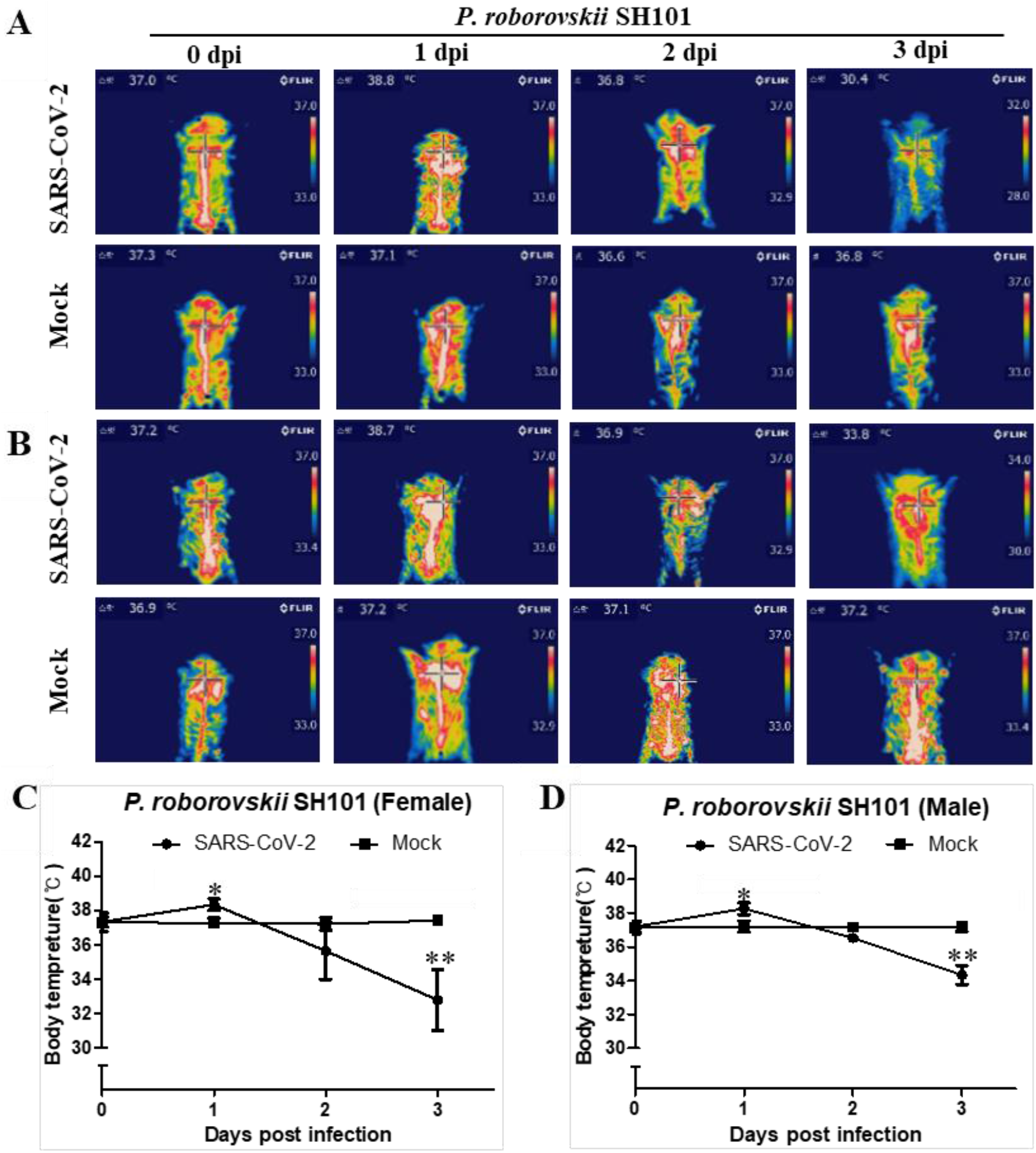
The body temperature changes of *P. roborovskii* SH101 infected with SARS-CoV-2. (A, B) The representative infrared thermographic images of the female (A) and male (B) *P. roborovskii* SH101 for 0, 1, 2, 3 days post-infection of SARS-CoV-2. (C, D) The body surface temperatures on the chest, as close as possible to the lung, of the female (C) and male (D) *P. roborovskii* SH101 hamster infected with SARS-CoV-2 (n = 6). The body temperatures were measured by selecting the highest temperature spot on the thermal images and are presented as mean ± SD. The statistical significances are marked on the graphs as * *P* < 0.05 and ** *P* < 0.01.

Another unusual feature of human COVID-19 is the widespread thrombosis of small and large vessels, contributing to morbidity and mortality^28–30^. It was also known that fibrinolysis was elevated in human COVID-19 due to systemic multiorgan thrombosis by SARS-CoV-2 infection [31,32]. The levels of fibrin degradation products, D-dimer and FDP, were elevated in the plasmas of the SH101 hamsters infected with SARS-CoV-2 (Fig 3A ~ D), suggesting the occurrence of systemic infection in the hamsters as in the case of human COVID-19.

**Fig 3.**
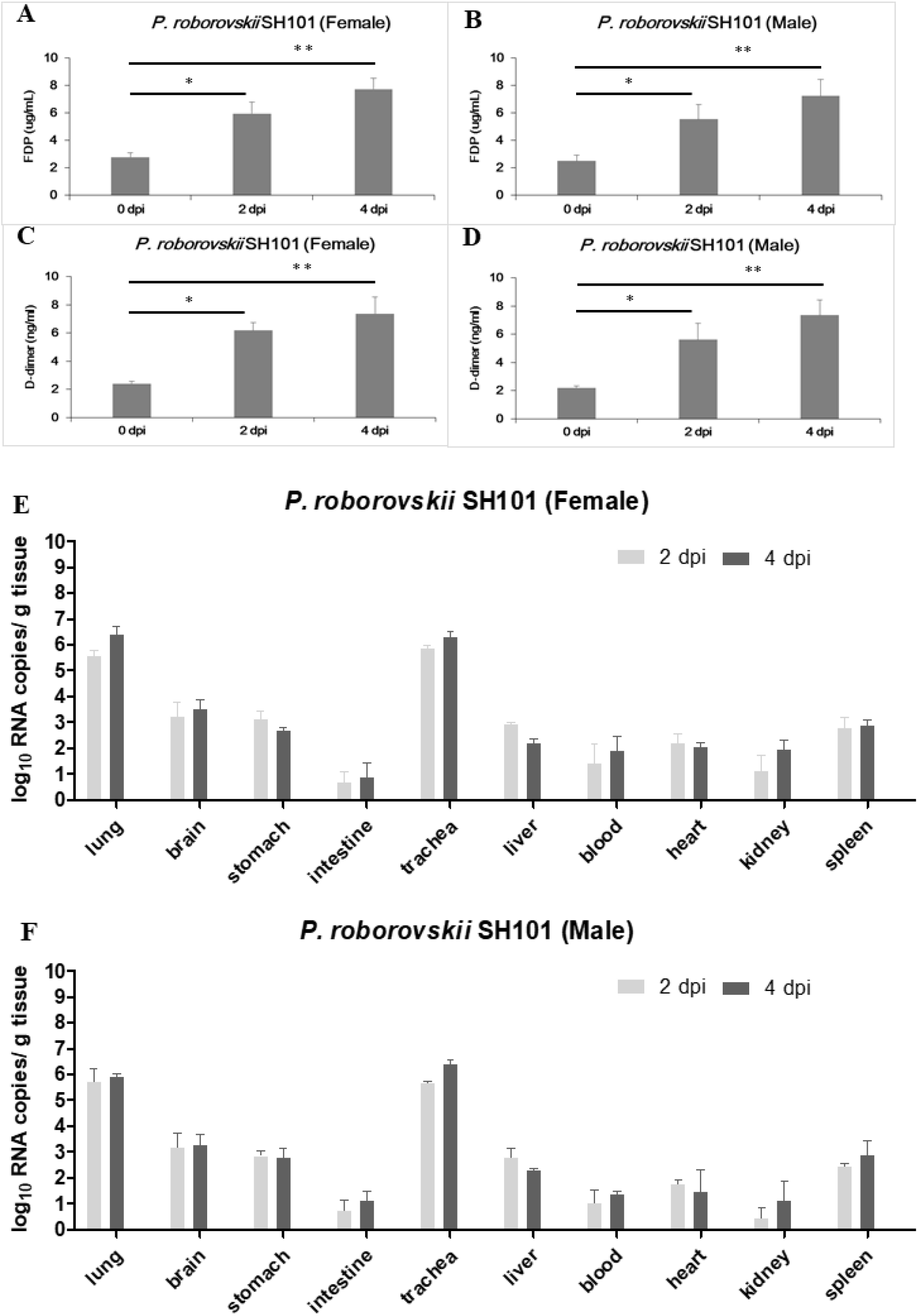
Thrombosis and viral replication in *P. roborovskii* SH101 infected with SARS-CoV-2. (A, B) The levels of fibrin degradation products (FDP) in the plasma of the female (A) and male (B) *P. roborovskii* SH101 hamsters at 2 and 4 dpi of SARS-CoV-2 (n = 3). (C, D) The levels of D-dimer in the plasma of the female (C) and male (D) *P. roborovskii* SH101 hamsters at 2 and 4 dpi of SARS-CoV-2 (n = 3). (E, F) The viral RNA levels in the lung, brain, stomach, intestine, trachea, liver, blood, heart, kidney, and spleen of the female (E) and male (F) *P. roborovskii* SH101 hamsters measured by RT-qPCR at 2 and 4 dpi of SARS-CoV-2. Data are presented as mean ± SD (n = 3). The statistical significances are marked on the graphs as * *P* < 0.05 and ** *P* < 0.01.

We next quantitated SARS-CoV-2 in the SH101 hamsters by quantitative RT-PCR after reverse transcription (RT–qPCR). The primary organs were collected from 3 randomly assigned individuals in each group at 2 and 4 dpi for analysis. High levels of viral RNA were detected in the homogenates of the lung and trachea, whereas lower levels were detected in the brain, stomach, intestine, liver, blood, heart, kidney, and spleen (Fig 3E, F). The detection of the viral RNAs in all organs from the SH101 hamsters confirmed serious systemic infection caused by SARS-CoV-2.

### The histological examinations of *P. roborovskii* SH101 infected with SARS-CoV-2 revealed severe inflammation in the lungs and minor pathologies in the liver and brain

After observing that *P. roborovskii* SH101 infected with SARS-CoV-2 closely represented the clinical manifestations of COVID-19 in humans, we examined the histopathological changes in each organ of the hamsters (Fig 4). Slides were carefully evaluated by a pathologist blinded to the treatment groups. As shown in Fig 4A, pathological examinations revealed severe inflammatory lesions in the lungs of the infected SH101 hamsters starting from 2 dpi. The lesions in the lungs were the typical features of severe viral pneumonia, as shown in diffuse alveolar damage (DAD) with hyaline membrane formation and type II pneumocyte desquamation at 2 dpi. The DAD pattern showed a mixture of exudative and proliferative features. The inflammatory lung lesions were extended across larger areas without any alveolar spaces due to the severity of inflammation at the last day, 4 dpi.

**Fig 4.**
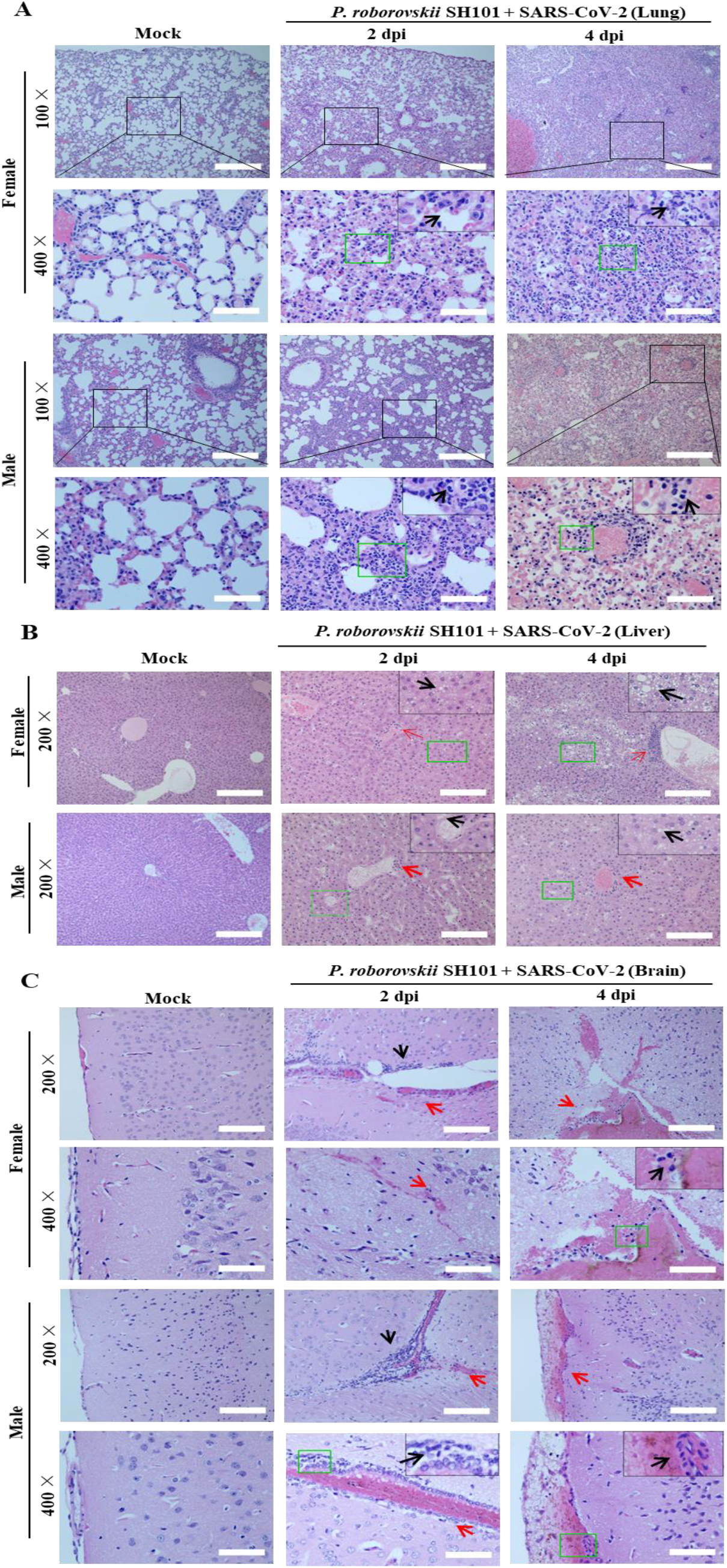
Histological examination of *P. roborovskii* SH101 infected with SARS-CoV-2. (A) The representative images of the H&E-stained histological sections of the lungs of *P. roborovskii* SH101 at 2 and 4 dpi of SARS-CoV-2. Multifocal interstitial pneumonia with thickened alveolar septa (green frame) and pinkish fibrinous infiltration of inflammatory cells (black arrows) are indicated. (B) The representative images of the H&E-stained histological sections of the livers of *P. roborovskii* SH101 at 2 and 4 dpi of SARS-CoV-2 showing pathologies. Focal and intraportal lymphoid cell aggregation and multifocal fatty changes are indicated by red and black arrows, respectively. (C) The representative images of the H&E-stained histological sections of the brains of *P. roborovskii* SH101 showing pathologies at 2 and 4 dpi. Subarachnoid hemorrhage and lymphocyte focal infiltration are indicated by red and black arrows, respectively. The scale bars represent 100 μm for 100 ×, 50 μm for 200 ×, and 20 μm for 400 ×.

In addition to the pulmonary injury, slight pathological damages were also observed in the livers and brains of the infected SH101 hamsters (Fig 4B). The pathological damages were observed in 4 out of the 6 livers at 2 dpi and 6 out of the 6 livers at 4 dpi, along with histological examinations. Multifocal fatty changes and portal lymphocytic infiltration observed in the livers are shown in Fig 4B. The hepatic injury also strongly supported that the SH101 hamsters infected with SARS-CoV-2 closely mimicked COVID-19 in human [33,34].

In brain specimens, like the liver, we also observed a focal infiltration of lymphocyte and/or subarachnoid hemorrhage in 4 out of the 6 brains at 2 dpi and 4 out of the 6 brains at 4 dpi (Fig 4C), suggesting possible neuroinflammation just like as in the case of COVID-19 disease [35,36]. Moreover, extravasation of red blood cells and hemosiderin pigments with focal infiltration of lymphocytes were also observed in the damaged brains. In terms of frequent neurological complications in COVID-19, the neurological damages (Fig 4C) in the infected SH101 hamsters also recapitulated the important clinical features of COVID-19. Despite high levels of SARS-CoV-2, the trachea was not damaged by SARS-CoV-2 infection, demonstrating another piece of similarity to COVID-19 [37]. The undamaged trachea despite the high viral titers implies that the trachea may function as an exhauster in the hamsters like the human counterpart. Other than these organs, no obvious histopathological changes were observed in the stomach, intestine, heart, kidney, and spleen (Fig S1).

### Comparison of *P. roborovskii* SH101 with the hACE2 transgenic mouse and Syrian hamster to study COVID-19

Among the various animal species investigated as an animal model for COVID-19, the hACE2 transgenic mice and Syrian hamsters are the most widely-used animal models to study human COVID-19. However, despite their popularity in COVID-19 studies, infections of the hACE2 transgenic mice or Syrian hamsters with SARS-CoV-2 resulted in infection mostly in the respiratory system. Considering their broad use in COVID-19 researches, these two animal models were analyzed in comparison with the SH101 hamster after SARS-CoV-2 infection. In agreement with previous studies, our experiments using hACE2 transgenic mice and Syrian hamsters confirmed their sensitivity to SARS-CoV-2 (Fig 5, 6 and Fig S2, S3) but failed to observe representative symptoms of COVID-19 such as fever induction, shaking chills, the right-over-left predominated pneumonia, *etc*.

**Fig 5.**
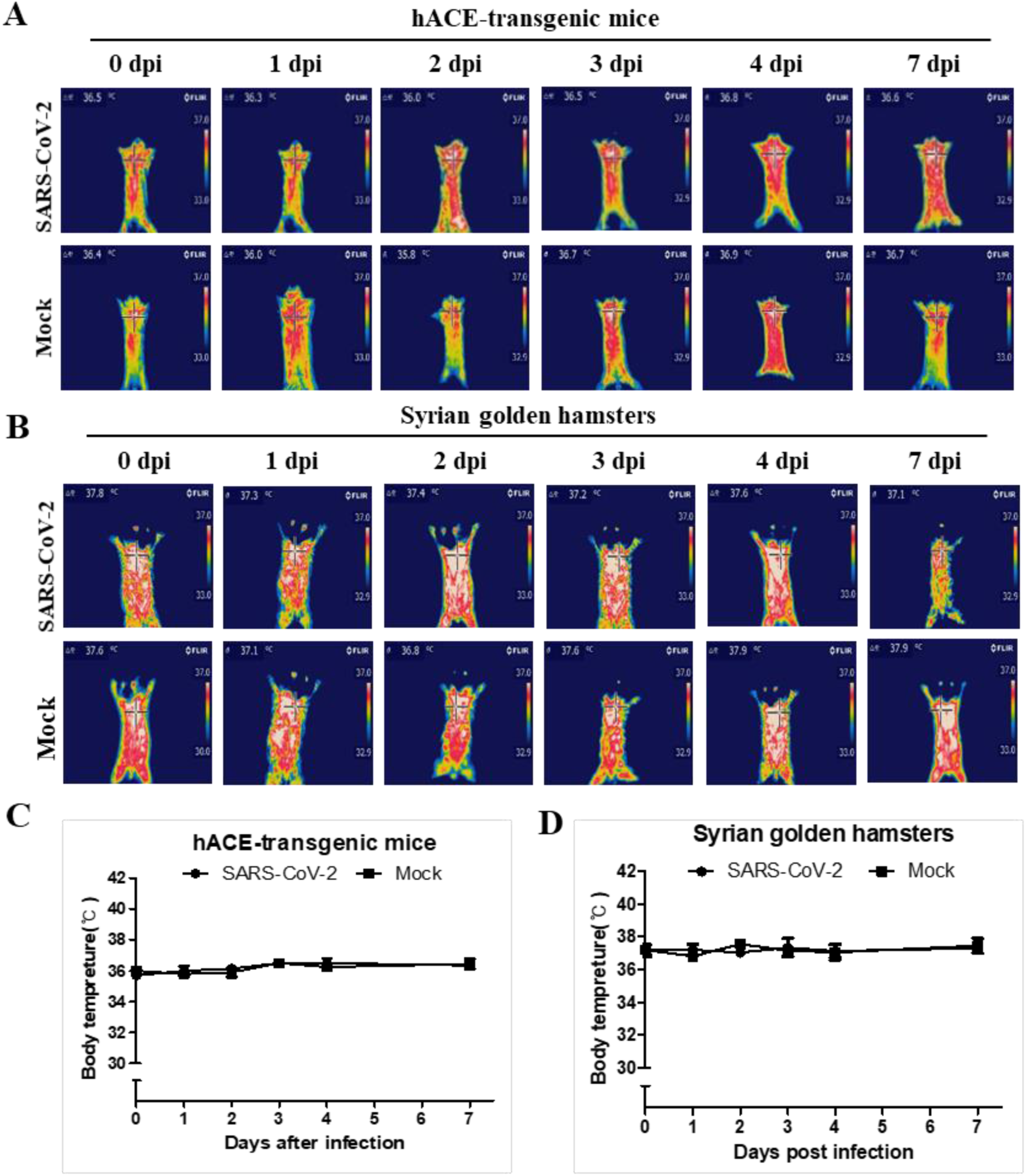
The body temperature changes of the hACE-transgenic mice and Syrian golden hamsters after infection of SARS-CoV-2. (A, B) The representative infrared thermographic images of the male hACE-transgenic mice (A) and male Syrian golden hamsters (B) at the indicated dpi of SARS-CoV-2. (C, D) The body temperatures on the chest, as close as possible to the lung, of the male hACE transgenic mice (C) and male Syrian golden hamster (D) at the indicated dpi (n = 6). The body temperatures were measured by selecting the highest temperature spot on the thermal images and are presented as mean ± SD.

**Fig 6.**
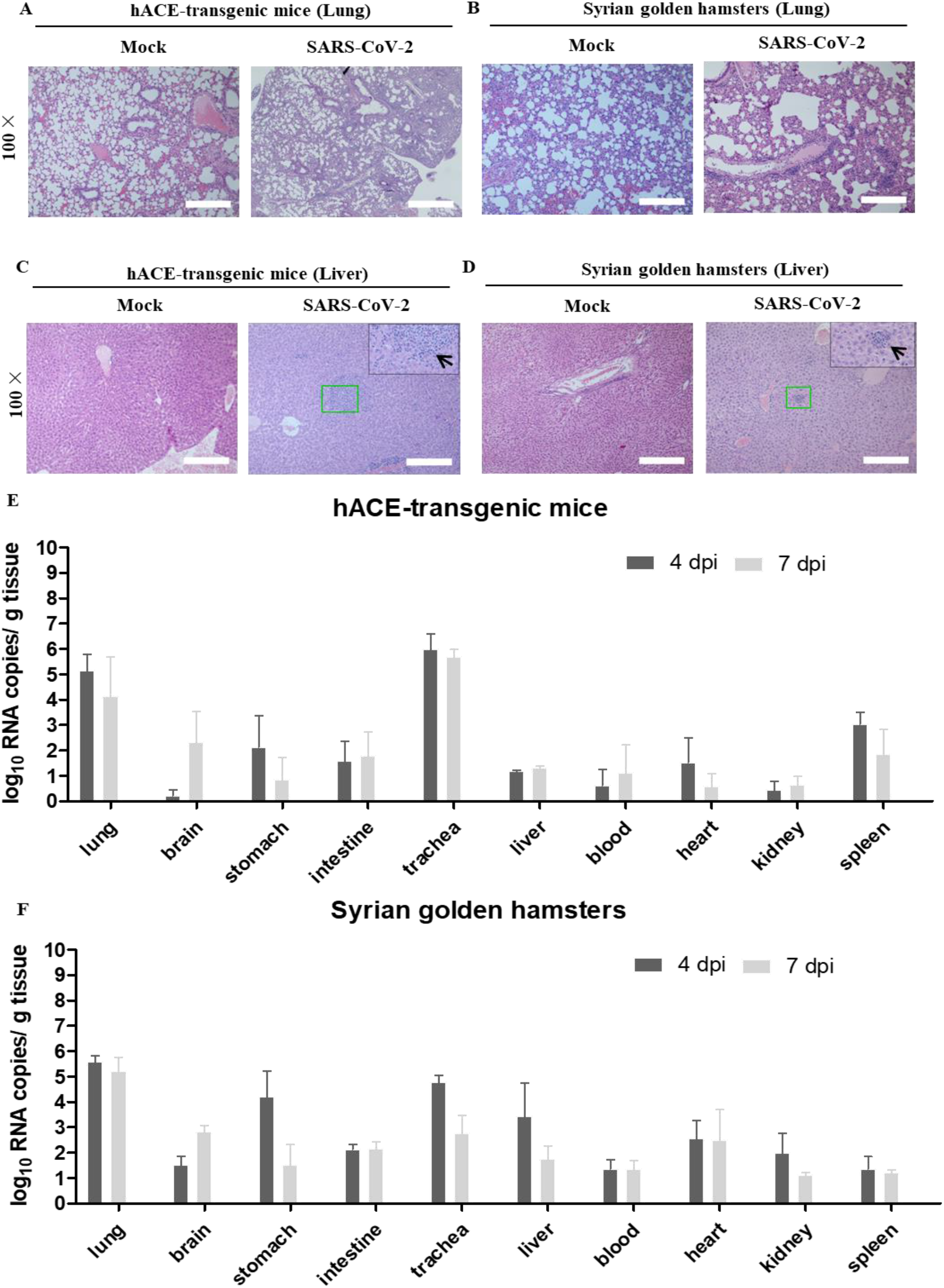
Histological examination and SARS-CoV-2 replication in the hACE-transgenic mice and Syrian golden hamsters infected with SARS-CoV-2. (A-D) The representative images of the H&E-stained histological sections of the lungs (A, B) and the livers (C, D) of the hACE-transgenic mice and Syrian golden hamsters at 7 dpi of SARS-CoV-2 showing pathologies. Multifocal interstitial pneumonia with thickened alveolar septa (green frame) and infiltration of inflammatory cells (black arrows) are indicated. (E, F) The viral RNA levels in the primary organs, lung, brain, stomach, intestine, trachea, liver, blood, heart, kidney, and spleen, of the male hACE-transgenic mice (E) and the male Syrian golden hamsters (F) measured by RT-qPCR at 4 and 7 dpi of SARS-CoV-2. Data are present as mean ± SD (n = 3). The scale bars represent 100 μm for 100 × magnifications.

The fever induction and shaking chills are the most common and primary symptoms of COVID-19 that can be observed even in mild cases [26,27]. For the first time, we were able to show the cardinal symptoms of COVID-19, fever induction, shaking chills, and respiratory problems, in a SARS-CoV-2-infected animal just like in the human counterpart (Fig 1 ~ 4, Fig S4 ~ S7 and Video S1~S4), but not in other models (Fig 5, 6).

Both hACE2 transgenic mice and Syrian hamster with SARS-CoV-2 infection demonstrated certain degree of organ damages, heavily damaged lungs and slightly damaged livers but no detectable pathology in the brains (Fig 6A ~ D and Fig S3). Interestingly, the viral RNA copies in the lungs of SH101, the hACE2 transgenic mice, and Syrian hamsters were not much different, 6.4 ± 0.4 (4 dpi), 5.7 ± 0.3 (4 dpi), and 5.6 ± 0.2 (4 dpi), respectively (Fig 6E, F). The dramatic differences of clinical manifestations between current models and SH101 despite of the similar range of virus titers (Fig 3E, F and Fig 6E, F) suggests that a physiological makeup of *P. roborovskii* SH101 would be different from those of the other two animals.

It has been well studied that young animals were significantly less sensitive to SARS-CoV-2 [38,39]. While hACE2 transgenic mice and Syrian hamsters displayed age-dependent sensitivity, SH101 hamsters as young as two-month-old were able to show serious symptoms by SARS-CoV-2 infection (Fig S4 ~ S7). The symptomatic severities of the young hamsters assessed by the organ damages were comparable to those of adults (Fig 4 and Fig S1, S5, S6). The virus titers of the young SH101 were 7.0 ± 0.3 in the lung at 4 dpi (Fig S7) like those of adults (Fig 3E, F). Despite the general similarity between the young and adult hamsters, SARS-CoV-2 infection led to a 100% mortality rate in the young SH101 hamsters by 4 dpi.

## Discussion

Multiple animal species have been investigated as an animal model for COVID-19 since the onset of the pandemic [38,39]. Among the developed animal models to study COVID-19, the hACE2 transgenic mice and Syrian hamsters are the most favorable animal models currently because of the merit of easy handling of small animals [13,15]. Despite of their popularity in the COVID-19 studies, the common use of these two animals for SARS-CoV-2 infection needs to be reconsidered due to their limits in the implementation of human COVID-19. This work confirmed that SARS-CoV-2 infections are mostly localized in the respiratory systems of the hACE2 transgenic mice and Syrian hamsters (Fig 5, 6 and Fig S2, S3) in agreement with previous studies [13,15]. COVID-19, however, is known to display a various degree of systemic infection in addition to localized respiratory infection, even in mild cases, indicating that the hACE2 transgenic mice and Syrian hamsters are not ideal models to study human COVID-19.

The repertory of clinical presentations of COVID-19 is far exceeding respiratory symptoms, including diarrhea, loss of sense of smell or taste, neuroinflammation manifested as encephalitis, meningitis, acute cerebrovascular disease, GBS, multisystem inflammatory syndrome, thrombosis, hyperfibrinolysis, *etc* [4,5]. Given the nature of the systemic infection of COVID-19, an ideal COVID-19 animal model should represent not only most respiratory infection but also other systemic infection observed in human COVID-19. Unfortunately, none of the current COVID-19 animal models reproduce the salient clinical and pathological features of human COVID-19^22^. Even primate models, the closest animal models to humans, did not reproduce the clinical symptoms from the systemic infection of human COVID-19 [16,38]. It is therefore pivotal to develop an animal model for the systemic infection replicating most clinical aspects of human COVID-19. In this regard, *P. roborovskii* SH101 would be an ideal animal model for evaluating antiviral therapeutic agents and vaccines, as well as understanding the pathogenesis of human COVID-19. Interestingly, *P. roborovskii* SH101 infected with SARS-CoV-2 recapitulated the very interesting COVID-19-specific facet, the right-predominated pneumonia (Fig 1E), which are uniquely observed in human COVID-19.

*P. roborovskii* SH101 not only reproduced the salient clinical and pathological features of human COVID-19 but also demonstrated unusual rapid progression once infected. The SARS-CoV-2-infected hamsters are observed with fever induction early on 1 ~ 2 dpi, followed by rapid progression into the terminal stage to death by 3 ~ 4 dpi (Fig 2). The unusual rapid progression in addition to clinical presentations would provide an ideal animal model to investigate disease mechanisms and to develop drugs and vaccines for COVID-19. These characteristics of the SH101 hamster model seem to eliminate the necessity for the primate models of COVID-19.

SARS-CoV-2 is an RNA virus that quickly evolves into various strains by rapid mutations. In fact, multiple strains of SARS-CoV-2 have already emerged [40], and thus the development of animal models recapitulating most of the clinical manifestations of COVID-19 would be of utmost importance not only for overcoming COVID-19 but also for understanding the pathogenesis. Further genome analyses of *P. roborovskii* SH101 in comparison with SARS-CoV-2-resistant hamsters or with SARS-CoV-2-susceptible hamsters for the local respiratory infection, such as Syrian hamsters, would provide critical clues for the understanding of human COVID-19.

## Materials and methods

### Ethics approval

All procedures involving the mice and hamsters were accredited with the approval of the Institutional Animal Care and Use Committee (IACUC), in compliance with the guidelines of the Ethics Committee of Jeonbuk National University Laboratory Animal Center Guidelines on the Care and Use of Animals for Scientific Purposes. All animal experiments in this study were in accordance with the ARRIVE guidelines and checklist. All the animals used in this research were handled in a manner consistent with CDC/ABSA/WHO guidelines for the prevention of human infection with the SARS-CoV-2 virus.

### Viruses and cells

The SARS-CoV-2 strain HB-01 was obtained from the National Culture Collection for Pathogens (NCCP) of the Korea Disease Control and Prevention Agency (KDCA). The complete genome for this SARS-CoV-2 has been submitted to GISAID (identifier: BetaCoV/Wuhan/IVDC-HB-01/2020|EPI_ISL_402119), and deposited in the China National Microbiological Data Center (accession number NMDC10013001 and genome accession number MDC60013002-01). Preparation of seed SARS-CoV-2 stocks and isolation of the virus were performed in Vero cells, which were maintained in Dulbecco’s modified Eagle’s medium (DMEM) supplemented with 10% fetal bovine serum (FBS), 100 IU/ml penicillin, and 100 μg/ml streptomycin, and incubated at 37°C, 5% CO_2_.

### Animal experiments

Six-week-old hACE2 transgenic mice (K18-hACE2 strain) having the human angiotensin I converting enzyme (peptidyl-dipeptidase A) 2 (hACE2) gene under the human cytokeratin 18 (K18) promoter [41] on chromosome 2 (99,209,508-99,220,724) and Syrian hamsters (*Mesocricetus auratus*) were purchase from Alpha biochemicals Co (http://alphabiochemicals.com). *P. roborovskii* SH101 was a laboratory inbred hamster strain maintained in the Jinis Biopharmaceuticals Inc. (Wanju, Jeonbuk, Republic of Korea). We deposited *P. roborovskii* SH101 for distribution in Alpha biochemicals Co.

For animal experiments, three animals of each animal species were housed in one cage and were provided with food (10% kcal as fat intake; D12450B; Research Diets Inc.) and water ad libitum. A 12-hour light-dark cycle was maintained (lights on at 8:30 PM daily) in a room of the ABL3 lab of Jeonbuk National University with controlled temperature (22°C ± 1°C) and humidity (55% ± 5%) for the animals. After 2 weeks of acclimation in the ABL3 lab, 50 μl of SARS-CoV-2 solution containing 10^5^ TCID_50_ or mock (DMEM) was inoculated intranasal by using a pipette. The food consumption of each mouse group was monitored daily. For blood profiling and behavioral tests, animals were randomly assigned to each treatment group. Particularly, we declare that blinding was employed during animal allocation and data collection.

### Body temperature measurement

The body temperatures of the animals were measured by a high-precision thermal photographic method [42,43]. Each of the whole animal bodies was photographed by FLIR thermal imaging camera, and the thermal images were analyzed by DirA Program (FLIR Tools) to measure the body temperature. The highest temperature spot on the chest, close to the site of the lung, was allocated in each mouse to determine the body temperature. All data are presented as the mean ± standard deviation and were compared using paired Student’s t-tests.

### D-dimer and Fibrin degradation products (FDPs) Analysis

The hypercoagulable state of the SARS-CoV-2-infected hamsters was analyzed daily by D-dimer and FDPs assays during the experimental period. Whole blood was collected in 1.5 ml Eppendorf micro-centrifuged tube by cardiac puncture. Blood serum was separated by centrifugation at 2,500 × g for 10 min and stored at 25°C until used. The D-dimer and FDPs concentrations of serum samples were determined by the double antibody sandwich method using mouse D-dimer and mouse FDPs ELISA kits (Sunlong Biotech, Hangzhou, Zhejiang, China) as described previously [44]. Briefly, 20 ~ 23 times diluted sera with the dilution buffer were used to quantitate D-dimer or FDPs in a micro-ELISA strip plate pre-coated with mouse anti-D-dimer or anti-FDPs monoclonal antibody. The concentrations of D-dimer and FDPs were calculated from the OD values of each sample using a standard curve.

### Preparation of total RNAs from the primary organs

After euthanized each mouse of the experimental groups, each mouse was dissected to isolate primary organs; lung, brain, liver, pancreas, kidney, stomach, trachea, spleen, small intestine, heart, and colon. Tissue homogenates (100 mg/ml) were prepared by homogenizing perfused organs using an electric homogenizer for 2 min 30 s in DMEM. The homogenates were centrifuged at 3,000 × rpm for 10 min at 4°C. The supernatants were collected and stored at −80°C until virus quantifications.

### Quantitation of SARS-CoV-2 titers

Total RNA was extracted from the supernatants of the organ homogenates using the RNeasy Mini Kit (QIAGEN, Hilden, Germany), and reverse transcription was performed using the PrimerScript RT Reagent Kit (TaKaRa, Japan) following the manufacturers’ instructions. RT–qPCR reactions were performed using the PowerUp SYBG Green Master Mix Kit (Applied Biosystems, Waltham, MA, USA), in which samples were processed in duplicate using the following cycling protocol: 50°C for 2 min, 95°C for 2 min, followed by 40 cycles at 95°C for 15 s and 60°C for 30 s, and then 95°C for 15 s, 60°C for 1 min, 95°C for 45 s. The primer sequences used for RT–qPCR is targeted against the envelope (E) gene of SARS-CoV-2 and are as follows: forward: 5′-GCCTCTTCTCGTTCCTCATCAC-3′, reverse: 5′-AGCAGCATCACCGCCATTG -3′. The PCR products were verified by sequencing using the dideoxy method on an ABI 3730 DNA sequencer (Applied Biosystems, Waltham, MA, USA). During the sequencing process, amplification was performed using specific primers. The sequencing reads obtained were compared with the NCBI database. The SYBR green real-time PCR standard curve was generated by serial ten-fold dilutions of recombinant plasmid with a known copy number (from 7 ×10^7^ to 7 × 10^1^ copies per μl). These dilutions were tested and used as quantification standards to construct the standard curve by plotting the plasmid copy number against the corresponding threshold cycle values (Ct). Results were expressed as log_10_-transformed numbers of genome equivalent copies per ml of sample. The Ct values of each sample were used to quantitate the virus titers by using the standard curve.

### Histology

About halves of the primary organs (lung, brain, liver, pancreas, kidney, stomach, trachea, spleen, small intestine, heart, and colon) were isolated from 3 adult postmortem *P. roborovskii* SH101 at 2 dpi (euthanized hamsters) and 4 dpi (hamster carcasses at terminal stage), respectively. The same primary organs were isolated for histological examination from 2-month-old *P. roborovskii* SH101 carcasses at 4 dpi, euthanized hACE2 transgenic mice at 7 dpi, and euthanized Syrian golden hamster at 7 dpi. Immediately after isolation, all organs were fixed by using 10% neutral-buffered formalin followed by embedded in paraffin and sectioned to be placed on glass microscopic slides (5 μm). After removing paraffin from the sections on the microscopic slides by hot water, the slides were air-dried and baked overnight at 65°C. The organ sections were stained with Hematoxyline and Eosin (H&E) with standard staining procedure [45,46]. The stained tissue images were observed in Apero ScanScope FL (Leica Biosystems, Germany).

### Statistical analysis

The statistical significance of SARS-CoV-2-infected and mock-treated samples was assessed by a one-way ANOVA multiple comparisons test. Statistical analyses were performed using GraphPad Prism 5 (GraphPad Software, La Jolla, CA, USA). A *p*-value less than 0.05 was considered statistically significant.

## Supporting Information

**S1 files. Figure S1~S7containing (PDF)**

**Fig S1:** The histological images of the primary organs of *P. roborovskii* SH101 post-infection of SARS-CoV-2. **Fig S2:** The body weight changes for the hACE-transgenic mice and Syrian golden hamsters post-infection of SARS-CoV-2. **Fig S3:** The histological examination results of the primary organs showing no pathological damages of the hACE-transgenic mice and Syrian golden hamsters post-infection of SARS-CoV-2. **Fig S4:** The changes of the body weight and the body temperatures of young *P. roborovskii* SH101 post-infection of SARS-CoV-2. **Fig S5:** The results of histological examination of the primary organs showing pathologies of young *P. roborovskii* SH101 post-infection of SARS-CoV-2. **Fig S6:** The histological examination results of the primary organs showing no pathological damages of young *P. roborovskii* SH101 post-infection of SARS-CoV-2. **Fig S7:** SARS-CoV-2 replication in young *P. roborovskii* SH101 after SARS-CoV-2 infection.

**S2 files. Video S1~S4 containing (ZIP)**

**Video S1:** General behavioral symptoms of male *P. roborovskii* SH101 infected with SARS-CoV-2 **Video S2:** The sneezing symptom of *P. roborovskii* SH101 infected with SARS-CoV-2. **Video S3:** The shaking chills of *P. roborovskii* SH101 infected with SARS-CoV-2. **Video S4:** General behavioral symptoms of young male *P. roborovskii* SH101 infected with SARS-CoV-2.

## Author Contributions

Conceptualization: Seong-Tshool Hong

Data curation: Chongkai Zhai, Mingda Wang

Formal analysis: Chongkai Zhai, Mingda Wang, Hea-Jong Chung, Md. Mehedi Hassan

Funding acquisition: Seong-Tshool Hong

Investigation: Seong-Tshool Hong

Methodology: Chongkai Zhai, Mingda Wang, Hea-Jong Chung, Md. Mehedi Hassan

Project administration: Chongkai Zhai, Mingda Wang

Resource: Chongkai Zhai, Mingda Wang, Hea-Jong Chung, Md. Mehedi Hassan

Software: Chongkai Zhai, Mingda Wang, Hea-Jong Chung, Md. Mehedi Hassan

Supervision: Seong-Tshool Hong

Validation: Chongkai Zhai, Mingda Wang

Visualization: Chongkai Zhai, Mingda Wang, Seungkoo Lee

Writing-original draft: Seong-Tshool Hong, Hyeon-Jin Kim

Writing-review & editing: Seong-Tshool Hong, Hea-Jong Chung

## Financial Disclosure Statement

This research was supported by JINIS BDRD Research Institute of JINIS Biopharmaceuticals Inc. (Initial of authors who received fund: S. H.). The funder had no role in study design, data collection and analysis, decision to publish, or preparation of the manuscript.

## Notes

### Competing Interest Statement

The authors have declared no competing interest.

