## supplementary Figures for "*Phodopus roborovskii* SH101 as a systemic infection model of SARS-CoV-2"

### Supporting Information

#### S1File: Fig S1~S7 containing

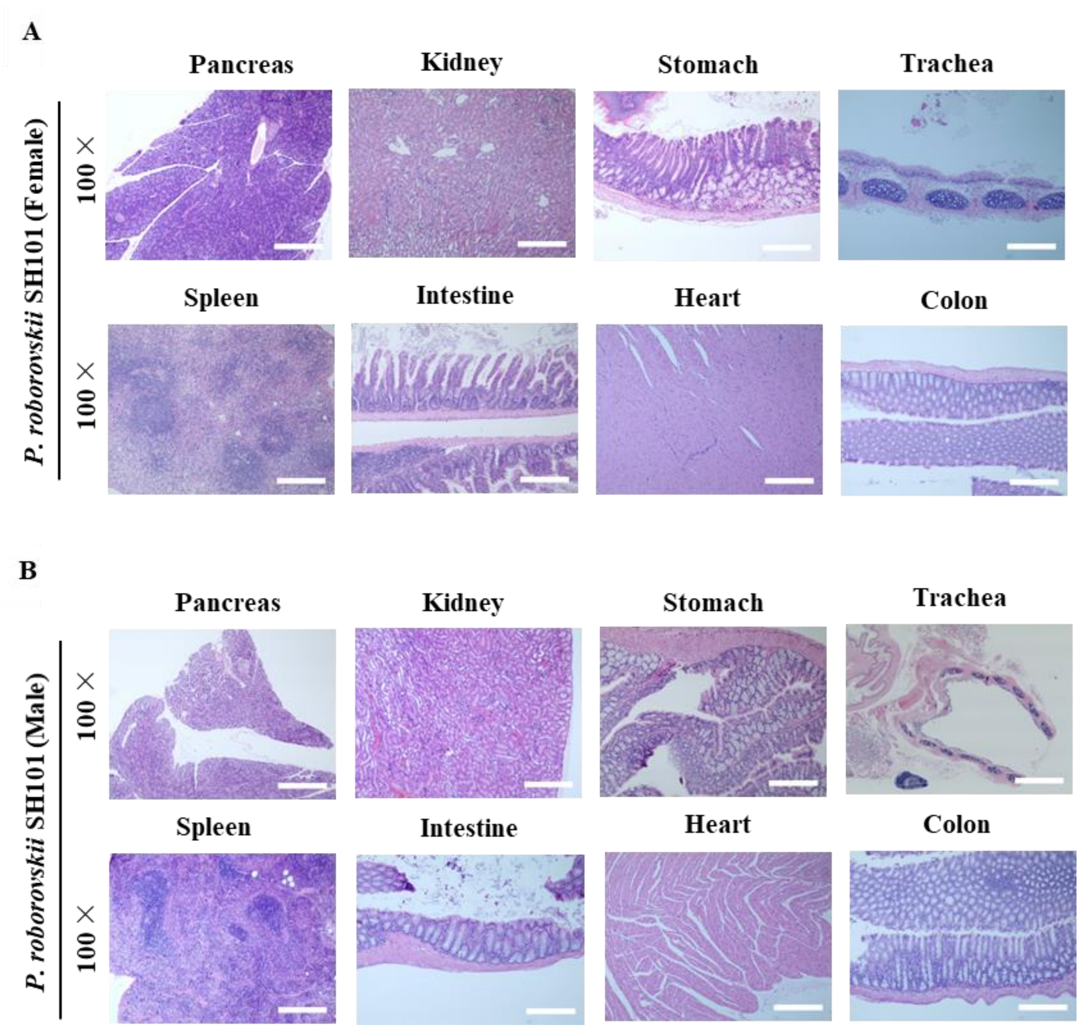


**Fig S1. The histological images of the primary organs of *P. roborovskii* SH101 infected with SARS-CoV-2.** The representative images of the H&E-stained histological sections without any pathologies in the primary organs of female (A) and male (B) *P. roborovskii* SH101 at 4 dpi of SARS-CoV-2. The scale bars represent 100 μm for 100 × magnifications.

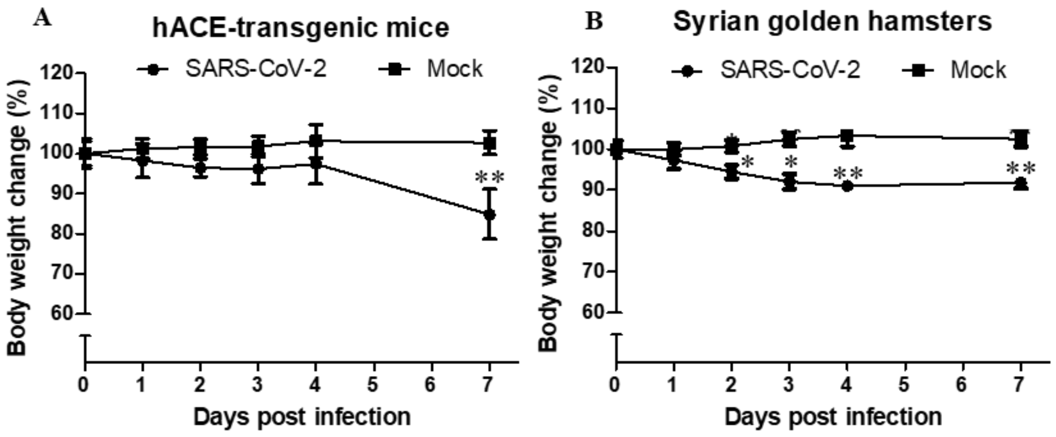


**Fig S2. The body weight changes for the hACE-transgenic mice and Syrian golden hamsters infected with SARS-CoV-2.** The body weights of the male hACE-transgenic mice (A**)** and the male Syrian golden hamsters (B) infected with SARS-CoV-2 were measured daily for 7 days (n = 6). Data are presented as mean ± SD. The statistical significances are marked on the graphs as * $P < 0.05$ and ** $P < 0.01$.

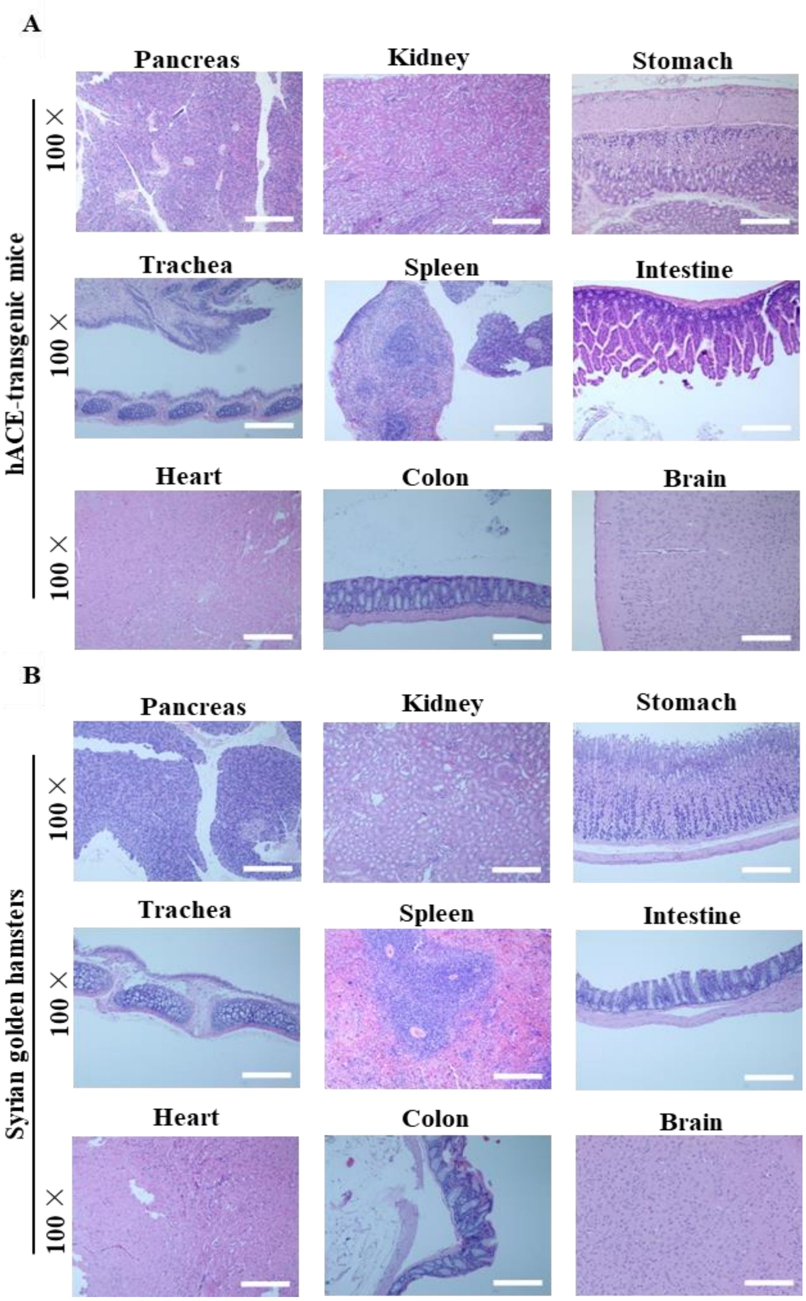


**Fig S3. The histological examination results of the primary organs showing no pathological damages of the hACE-transgenic mice and Syrian golden hamsters infected with SARS-CoV-2.** (A, B) The representative images of the H&E-stained histological sections of the primary organs of the male hACE-transgenic mice (A) and male Syrian golden hamsters (B) at 7 dpi of SARS-CoV-2. The scale bars represent 100 μm for 100 × magnification.

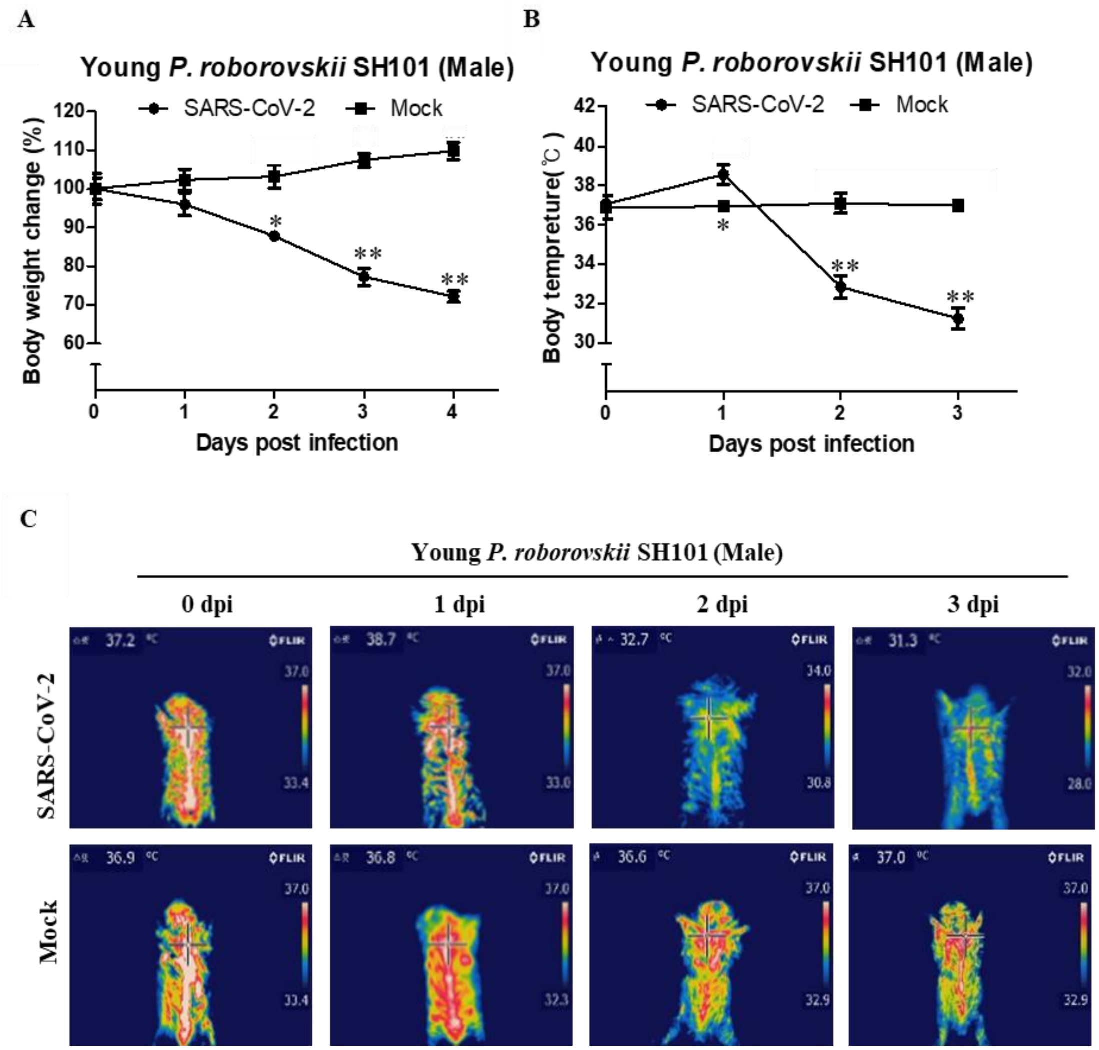


**Fig S4. The changes of the body weight and the body temperatures of young *P. roborovskii* SH101 post-infection of SARS-CoV-2.** (A) The body weights of the young male *P. roborovskii* SH101 post-infection of SARS-CoV-2 were measured daily for 4 days (n=3). (B) The surface body temperatures of the young male *P. roborovskii* SH101 post-infection of SARS-CoV-2 were measured daily for 4 days (n = 3). (C) The representative infrared thermographic images of the young male *P. roborovskii* SH101 post-infection of SARS-CoV-2. The surface body temperatures were measured by selecting the highest temperature spot on the thermal images where the lung is located, and are presented as mean ± SD. The statistical significances are marked on the graphs as * $P < 0.05$ and ** $P < 0.01$.

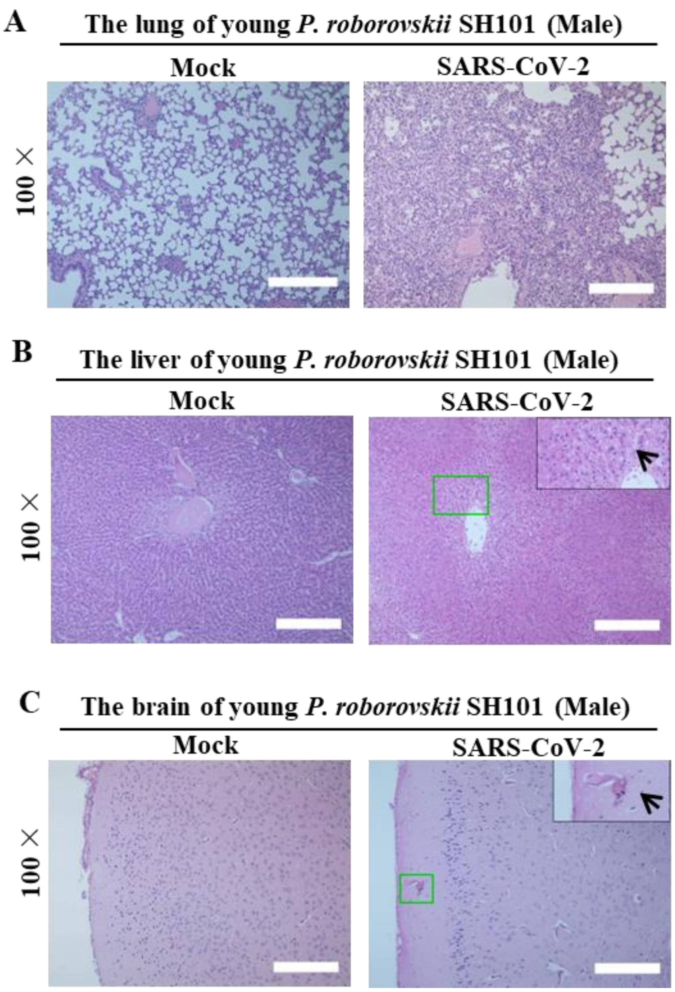


**Fig S5. The results of histological examination of the primary organs showing pathologies of young *P. roborovskii* SH101 post-infection of SARS-CoV-2. a-c,** The representative images of the H&E-stained histological sections of the lungs (a), brains (b), and livers (c) of the young *P. roborovskii* SH101 at 4 dpi. Multifocal interstitial pneumonia with thickened alveolar septa (green frame) and pinkish fibrinous infiltration of inflammatory cells (black arrows) can be observed. The scale bars represent 100 μm for 100 × magnification.

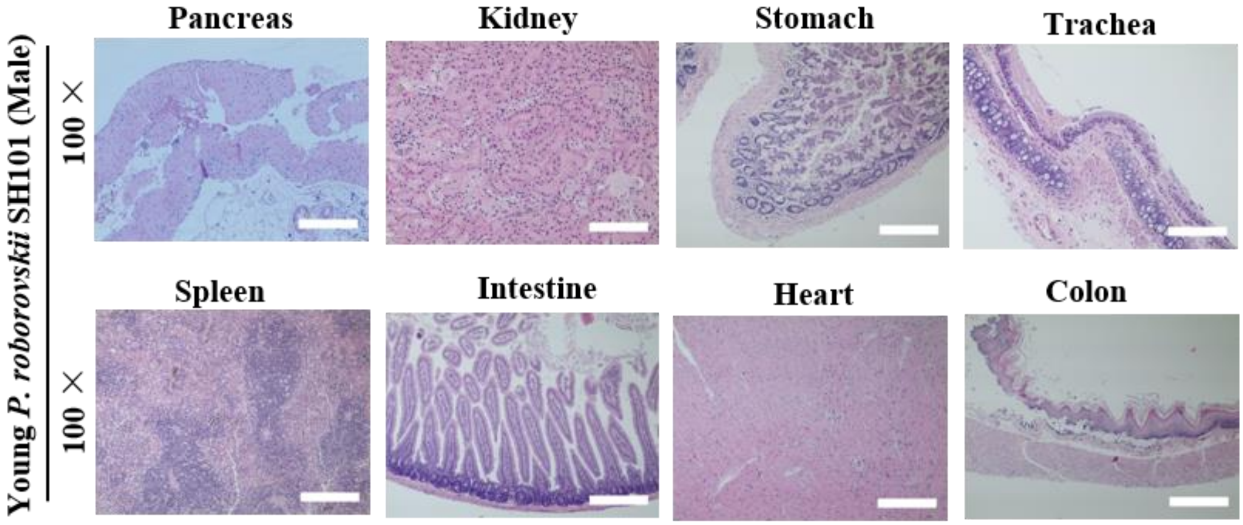


**Fig S6. The histological examination results of the primary organs showing no pathological damages of young *P. roborovskii* SH101 infected with SARS-CoV-2.** The representative images of the H&E-stained histological sections of the undamaged primary organs of the young *P. roborovskii* SH101 infected with SARS-CoV-2 at 4 dpi are shown. The scale bars represent 100 μm for 100 × magnification.

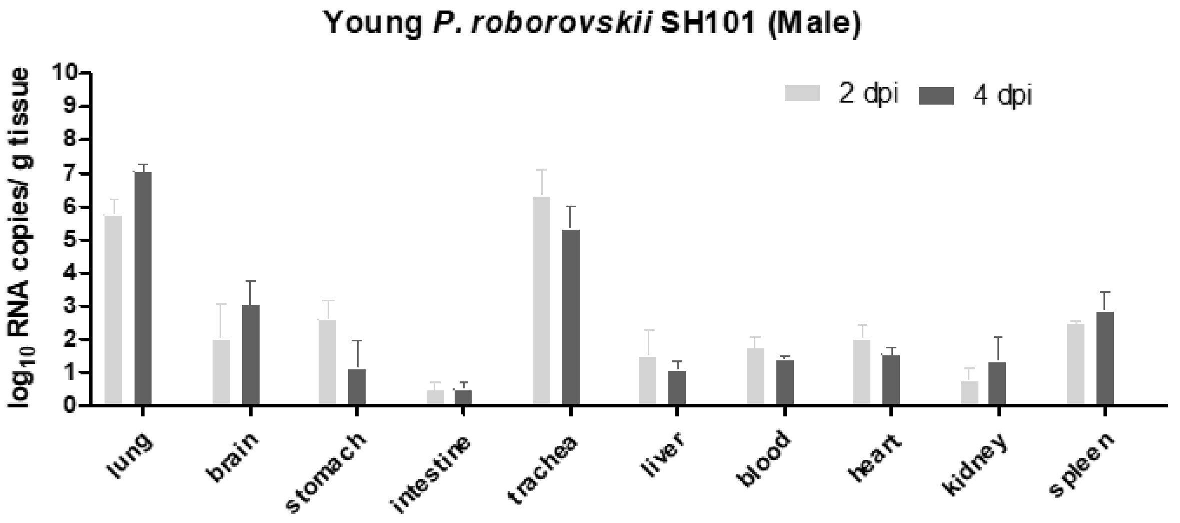


**Fig S7. SARS-CoV-2 replication in young *P. roborovskii* SH101 after SARS-CoV-2 infection.** The viral RNA levels in the lung, brain, stomach, intestine, trachea, liver, blood, heart, kidney, and spleen of the young *P. roborovskii* SH101 were measured by RT-qPCR 2 and 4 dpi of SARS-CoV-2 (n = 3). Data are present as mean ± SD.

**S2 File: Video S1~S4 containing**

**Video S1:** General behavioral symptoms of male *P. roborovskii* SH101 infected with SARS-CoV-2.

**Video S2:** The sneezing symptom of *P. roborovskii* SH101 infected with SARS-CoV-2.

**Video S3:** The shaking chills of *P. roborovskii* SH101 infected with SARS-CoV-2.

**Video S4:** General behavioral symptoms of young male *P. roborovskii* SH101 infected with SARS-CoV-2.
